# Phosphorylation of STIM1 at ERK/CDK sites is dispensable for cell migration and ER partitioning in mitosis

**DOI:** 10.1101/2021.07.06.451280

**Authors:** Ayat S. Hammad, Fang Yu, Welathanthrige S. Botheju, Asha Elmi, Ethel Alcantara-Adap, Khaled Machaca

**Author notes:** Correspondence: Khaled Machaca, Weill Cornell Medicine Qatar, Education City - Qatar Foundation– Doha, Qatar. Indicates equal contribution (Ayat S. Hammad, Fang Yu).

## Abstract

Store-operated Ca^2+^ entry (SOCE) is a ubiquitous Ca^2+^ influx required for multiple physiological functions including cell motility. SOCE is activated in response to depletion of intracellular Ca^2+^ stores following the activation of endoplasmic reticulum (ER) Ca^2+^ sensor STIM1 which recruits the plasma membrane (PM) Ca^2+^ channel Orai1 at ER-PM junctions to induce Ca^2+^ influx. STIM1 is phosphorylated dynamically and this phosphorylation has been implicated in several processes including SOCE inactivation during M-phase, maximal SOCE activation, ER segregation during mitosis, and cell migration. Human STIM1 has 10 Ser/Thr residues in its cytosolic domain that match the ERK/CDK consensus phosphorylation. We recently generated a mouse knock-in line where wild-type STIM1 was replaced by a non-phosphorylatable STIM1 with all 10 S/T mutated to Ala (STIM1-10A). Here, we generate mouse embryonic fibroblasts (MEF) the STIM1-10A mouse line and a control MEF line (WT) that express wild-type STIM1 from a congenic mouse strain. These lines offer a unique model to address the role of STIM1 phosphorylation at endogenous expression and modulation levels in contrast to previous studies that relied mostly on overexpression. We show that STIM1 phosphorylation at ERK/CDK sites is not required for SOCE activation, cell migration, or ER partitioning during mitosis. These results rule out STIM1 phosphorylation as a regulator of SOCE, migration and ER distribution in mitosis.

## INTRODUCTION

Ca^2+^ is a ubiquitous second messenger engaged downstream of agonist-dependent receptor stimulation that leads to Ca^2+^ release from intracellular ER stores, and is typically followed by Ca^2+^ influx from the extracellular space through store-operated calcium entry (SOCE). SOCE is a prominent Ca^2+^ influx pathway that contributes to many cellular and physiological functions (Lacruz and Feske, 2015; Prakriya and Lewis, 2015; Putney, 1986). Ca^2+^ store depletion causes STIM1, an ER Ca^2+^ sensor, to oligomerize, leading to its accumulation at endoplasmic reticulum (ER) and plasma membrane (PM) junctions, where it binds to Orai1, the pore-forming subunit of the Ca^2+^ release-activated Ca^2+^ (CRAC) channel, activating local Ca^2+^ entry.

STIM1 is a single pass protein that localizes to the ER membrane with luminal EF-hand domains that sense store Ca^2+^ levels and support STIM1 oligomerization in response to store depletion (Fig. 1A) (Prakriya and Lewis, 2015). The STIM1 cytoplasmic domain contains three coiled-coil region that regulate STIM1 activation and binding to Orai1 (see Fig. 1A). The C-terminal end of the molecule contains a Pro/Ser rich domain (S/P) where many phosphorylated residues are concentrated. An EB1 binding domain (EB) that links STIM1 and by extension the ER to microtubules (MT). As well as a polybasic domain (PB) that mediates interactions with negative lipid headgroups at the PM to support the localization of activated STIM1 to ER-PM junctions.

**Figure 1.**
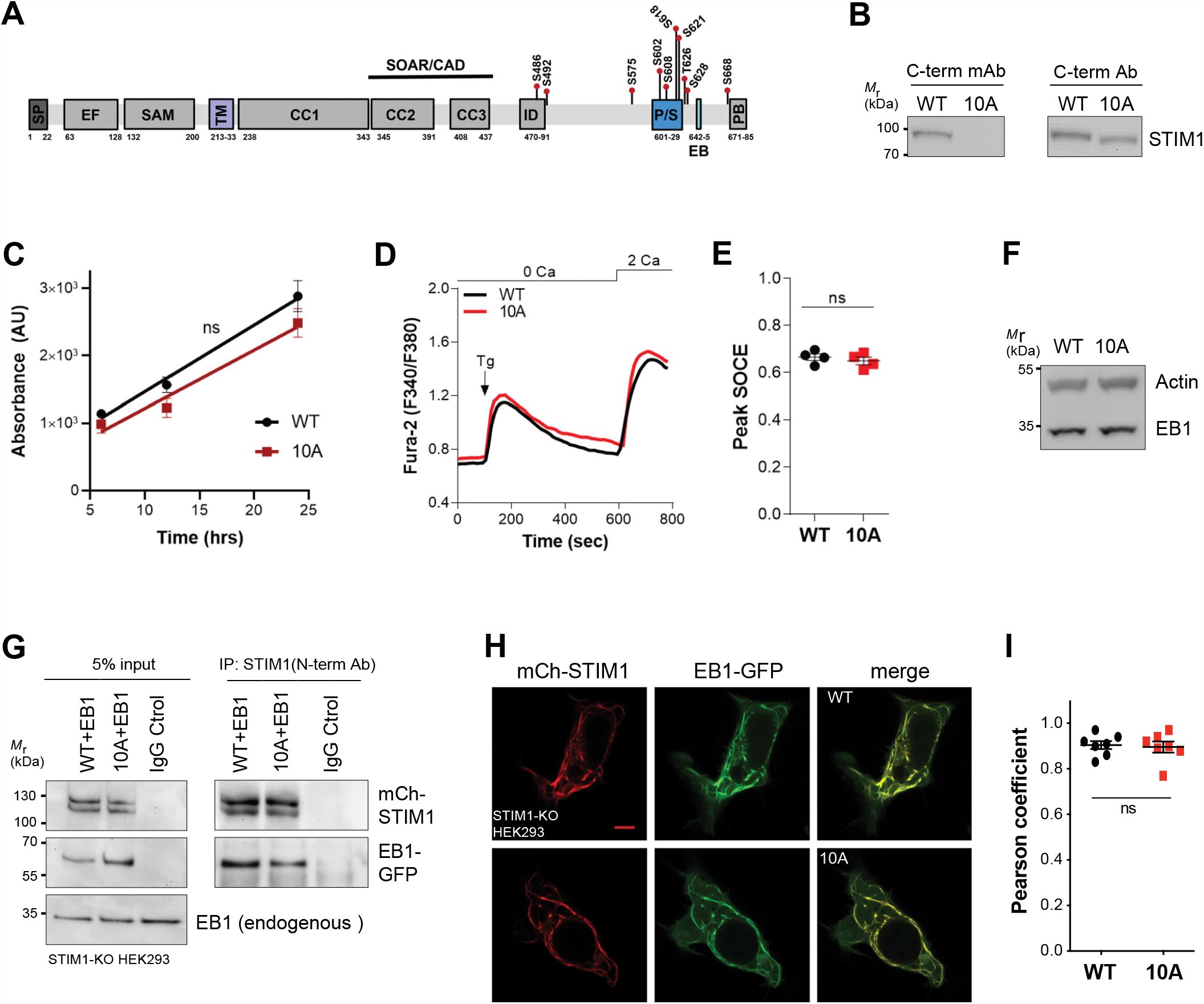
STIM1 phosphorylation state does not affect cell proliferation or SOCE activation. **(A)** Cartoon of STIM1 domains with the numbering below based on human STIM1 sequence (Q13586). SP: signal peptide; EF: EF-hand motif; SAM: SAM domain; TM: transmembrane domain; CC1, CC2, CC3: coiled coils; SOAR/CAD: Orai1 activating region that directly binds Orai1; ID: inactivation domain; P/S: Pro/Ser rich region; EB: EB1 binding ‘TRIP’ domain; PB: poly-basic domain. **(B)** Genotyping of STIM1 in MEFs generated from WT and STIM1-10A mice by Western blot with two different antibodies. Images represent two independent experiments. **(C)** Proliferation of the WT and 10A MEF cell lines at 6, 12 and 24 hrs (n = 5 independent experiments, mean ± SEM; ns, not significant, paired t test). **(D-E)** Representative traces (D) and summary data of SOCE (E) in WT and STIM1-10A MEFs (n = 4 independent experiments, mean ± SEM; ns, not significant, paired t test). **(F)** Western blot to evaluate EB1 expression in MEFs. **(G)** Lysates of STIM1-KO HEK-293 cells co-expressing EB1-GFP with either mCherry-STIM1 WT or 10A mutant were subjected to immunoprecipitation with STIM1 N-terminal Ab followed by Western blot with antibodies to detect EB1 and STIM1. 5% of the input lysate is shown in the left panels. Images represent two independent experiments. **(H)** Colocalization of EB1-GFP with mCherry-STIM1 WT or 10A mutant in STIM1-KO HEK-293 cells. Representative Airyscan confocal images (scale bar, 5 μm). **(I)** Pearson correlation coefficient between mCh-STIM1 and EB1-GFP (n=7 cells from two independent experiments, mean ± SEM; ns, not significant, unpaired t test).

There is significant interest in STIM1 phosphorylation and its role in regulating STIM1 functions. STIM1 was originally isolated as a phosphoprotein at rest that is primarily phosphorylated on Ser residues with some Tyr phosphorylation detected (Manji et al., 2000). However, the physiological role of STIM1 phosphorylation remains vague and it is not clear how and if increased phosphorylation above basal levels regulate STIM1 function.

SOCE is known to inactivate during M-phase of the cell cycle (mitosis and meiosis), which correlates with STIM1 hyper-phosphorylation (Arredouani et al., 2010; Smyth et al., 2009; Yu et al., 2009). However, the STIM1 phosphorylation status has no impact on SOCE inactivation during meiosis (Yu et al., 2009). In contrast, it was proposed that STIM1 phosphorylation mediates SOCE inhibition in mitosis based mostly on overexpression studies (Smyth et al., 2009). Recently however it was conclusively shown that STIM1 phosphorylation does not inactivate SOCE in mitosis using a knock-in mouse that expresses only the non-phosphorylated form of STIM1 at native endogenous levels (Yu et al., 2019). It was rather proposed that SOCE inhibition during mitosis is due to remodeling of ER-PM junctions that prevents direct STIM1-Orai1 interactions (Yu et al., 2019).

In addition, STIM1 phosphorylation has been implicated in maximal SOCE activation and Orai1 recruitment (Pozo-Guisado et al., 2010; Yazbeck et al., 2017). STIM1 associates with microtubules (MT) through binding to the microtubule-plus-end-tracking protein EB1 (Grigoriev et al., 2008). EB1 binds STIM1 through the ‘TRIP’ sequence within the EB domain (Fig. 1A) that matches the SxIP EB1 binding consensus (Honnappa et al., 2009). This binding is disrupted following STIM1 phosphorylation presumably because the phosphorylated residues are in close proximity to the EB-domain (Pozo-Guisado et al., 2013; Smyth et al., 2012). The phosphorylation-dependent dissociation of STIM1 from EB1 has been postulated to be important for SOCE activation to release STIM1 from MT and allow its interaction with Orai1 (Pozo-Guisado et al., 2013). Furthermore, the phosphorylation driven dissociation of STIM1-EB1 has been argued to prevent mis-localization of the ER to the spindle during mitosis and its mis-partitioning (Smyth et al., 2012).

Finally, STIM1 phosphorylation has been shown to modulate cell migration. Cell migration requires fine spatial and temporal regulation of the actin cytoskeleton as well as Ca^2+^ signaling pathways, both of which are polarized in migrating cells (Hammad and Machaca, 2021). Ca^2+^ signaling is involved in the remodeling of cortical actin that underlies the formation of membrane protrusions such as lamelipodia, filopodia, and invadopodia (Prevarskaya et al., 2011; Ridley, 2011; Ridley et al., 2003). SOCE plays important roles in cell migration by regulating focal adhesions (FA) turnover differentially at the front and rear of migrating cells (D’Souza et al., 2020; Hammad and Machaca, 2021; Prevarskaya et al., 2011; Tsai et al., 2014; Yang et al., 2009). SOCE enhances FA and thus adhesion to the extracellular matrix (ECM) at the leading edge (Tsai et al., 2014), but also paradoxically SOCE is required for disassembly of FA at the trailing end (D’Souza et al., 2020). Consistently, modulating the STIM and/or Orai expression levels affects cell migration, including metastasis of cancerous cells (Chen et al., 2011; Chen et al., 2016a; Hammad and Machaca, 2021; Sun et al., 2014; Yang et al., 2009).

Overexpression of a STIM1 mutant where some of the ERK phosphorylation sites (S575, S608, and S621) were mutated to alanines reduced cell migration (Casas-Rua et al., 2015). Furthermore, phosphorylated STIM1 as detected by phospho-specific antibodies is enriched at the leading edge and membrane ruffles in migrating cells (Lopez-Guerrero et al., 2017). Both STIM1 and Orai1 have been proposed to interact with cortactin (CTTN), a major player in actin cytoskeleton remodeling (Lopez-Guerrero et al., 2020; Lopez-Guerrero et al., 2017). Phosphorylation of STIM1 to spatially regulate SOCE would be an attractive regulatory approach as it is dynamic and can be readily controlled spatially.

Most of the studies addressing the role of STIM1 phosphorylation relied on overexpression as it is difficult to modulate the phosphorylation status of endogenous STIM1. The 10 S/T residues that match the ERK/CDK phosphorylation consensus within the STIM1 cytosolic domain are highly conserved between human and mouse. Therefore, to further explore the role of STIM1 phosphorylation in a more physiological context, we generated a STIM1-10A knock-in mouse line (STIM1-10A), in which all 10 S/T residues were replaced by Ala (A) (Yu et al., 2019), and derived a MEF line from this mouse strain. This allowed us to test the role of phosphorylation of endogenous STIM1 on SOCE, cell migration and ER distribution during mitosis. We show that STIM1 phosphorylation does not affect SOCE activation or ER localization during mitosis. We further show that STIM1 phosphorylation plays a minor role in modulating the speed of individual cell migration, an effect that does not translate into a detectable phenotype in cell-sheet migration, cell invasion, or in the context of whole organismal development.

## RESULTS

### STIM1 phosphorylation is dispensable for SOCE and for cell proliferation

The C-terminal cytoplasmic domain of STIM1 contains 10 residues that match the minimal ERK/CDK consensus phosphorylation sequence ‘S/T-P’ that are mostly clustered around the P/S and EB domains (Fig. 1A). We had previously generated a knock-in mouse line that expresses a STIM1 isoform where all these residues are mutated to Ala and are as such non-phosphorylatable (Yu et al., 2019). We refer to this STIM1 mutant as 10A or non-phosphorylatable for simplicity here, although we did not alter potential Tyr phosphorylation sites in the cytoplasmic domain or potential luminal phosphorylation sites (Pollak et al., 2018; Yazbeck et al., 2017).

STIM1 phosphorylation at ERK sites, specifically S575, S608, and S621, has been argued to be required for SOCE activation (Pozo-Guisado et al., 2010). Therefore, to further investigate the role of STIM1 phosphorylation in SOCE activation we generated MEF cell lines from the STIM1-10A strain as well as from a congenic strain that expresses wild-type STIM1. As shown previously the STIM1-10A strain develops and reproduces normally without any overt defects (Fig. S1A). The genotype of immortalized MEFs was confirmed by PCR (Fig. S1B) and by Western blot using two different STIM1 antibodies (Fig. 1B) validated previously (Yu et al., 2019). The polyclonal antibody against the C-terminal region of STIM1 detects endogenous STIM1 in both WT and 10A MEFs (Fig. 1B). In contrast, a C-terminal monoclonal antibody raised against a region around P622, which is enriched with potential phosphorylation sites (Fig. 1A), recognizes WT but not 10A STIM1 (Fig. 1B). These data show that 10A MEFs express only the non-phosphorylatable STIM1, whereas WT MEFs express only WT STIM1.

Because SOCE has been implicated in and is modulated during cell cycle progression (Chen et al., 2016b; Machaca and Haun, 2000; Preston et al., 1991; Tani et al., 2007), we tested the proliferation rate of both MEF lines over a 24 hrs time period. Both lines proliferate with similar rates (Fig. 1C), arguing against a role for STIM1 phosphorylation in cell proliferation.

We next measured SOCE using the standard Ca^2+^ re-addition assay (Fig. 1D) and observed no differences in SOCE levels (Fig. 1E) or slope (Fig. S1C) between WT and 10A MEFs. These data are consistent with our previous findings on primary T cells from STIM1-10A and WT mice, which showed no differences in SOCE (Yu et al., 2019). We further do not observe any difference in Ca^2+^ store content assessed using thapsigargin release between the two cell lines (Fig. S1D). These results argue that STIM1 phosphorylation at ERK/CDK sites is not required for maximal SOCE activation.

### Non-phosphorylatable STIM1 interacts normally with EB1

As discussed above STIM1 phosphorylation has been shown to inhibit STIM1-EB1 interactions with implications on both SOCE activation and ER distribution during mitosis (Pozo-Guisado et al., 2013; Smyth et al., 2012). The EB1-binding motif ‘TRIP’ locates to the C-terminus of STIM1 near the phosphorylation sites (Fig. 1A, EB). We first confirmed that the expression levels of endogenous EB1 is similar in both the WT and STIM1-10A MEFs (Fig. 1F). To rule out the possibility that the 10A mutations alter STIM1 binding to EB1, we examined EB1 and STIM1 interaction by immunoprecipitation (IP) using a STIM1 N-terminal polyclonal antibody. STIM1-KO HEK293 cells (Emrich et al., 2019) were used to co-express EB1-GFP with either mCherry-tagged human STIM1-WT or -10A mutant (Yu et al., 2019). EB1 was pulled down with similar efficiency from cells expressing either WT or STIM1-10A (Fig. 1G, right panels). Furthermore, the IP input gels show that endogenous EB1, overexpressed EB1-GFP, and mCherry-STIM1 WT or 10A were expressed at similar levels (Fig. 1G, left). To test for STIM1-EB1 interactions in situ, we co-expressed EB1-GFP with either mCherry-STIM1 WT or -10A and show that both WT and STIM1-10A colocalized to similar levels with EB1 (Fig. 1H) as assessed by Pearson’s colocalization coefficient (Fig. 1I).

### STIM1 phosphorylation does not affect ER distribution during mitosis

STIM1 phosphorylation has been argued to be a regulatory mechanism that excludes ER from the mitotic spindle (Smyth et al., 2012). Normally, STIM1 dissociates from EB1 during mitosis, however, the non-phosphorylatable STIM1-10A was shown to associate with EB1 resulting ER mis-localization to the mitotic spindle (Smyth et al., 2012). Surprisingly though this was not associated with any mitotic defects in either Hela or HEK293 cells (Smyth et al., 2012). We were therefore interested in testing whether these observations hold in cells where endogenous STIM1 is non-phosphorylatable as afforded by the STIM1-10A MEF line.

To assess the distribution of the ER and STIM1 in mitotic cells we stained for endogenous STIM1 in naturally occurring mitotic WT or 10A MEFs (Fig. 2A). We avoided the use of nocodazole to enrich for cells in mitosis as this may be associated with partial mitotic arrest. STIM1/ER mis-localization was assessed by quantifying the % STIM1 that localizes to the DNA area stained by DAPI and showed no differences between the two cells lines (Fig. 2A), arguing that STIM1-10A at endogenous expression levels does not result in ER mis-localization to the spindle during mitosis. The results with STIM1 were confirmed using both calreticulin (Fig. 2C) and SERCA (Fig. S2A) as independent resident ER proteins, or by expressing mCherry-KDEL to label the ER (Fig. 2B). For the calreticulin experiments we stained the cells for tubulin to mark the spindle and observe no mis-localization of the ER to the spindle or DNA areas in the 10A cells (Fig. 2C). We further tested ER segregation in mitosis in *ex vivo* cultured bone marrow derived macrophages (BMDM) from WT and STIM1-10A mice by immunostaining (Fig. S2B) and observed no abnormality in ER localization in mitotic BMDM from 10A mice compared to WT mice (Fig. S2B).

**Figure 2.**
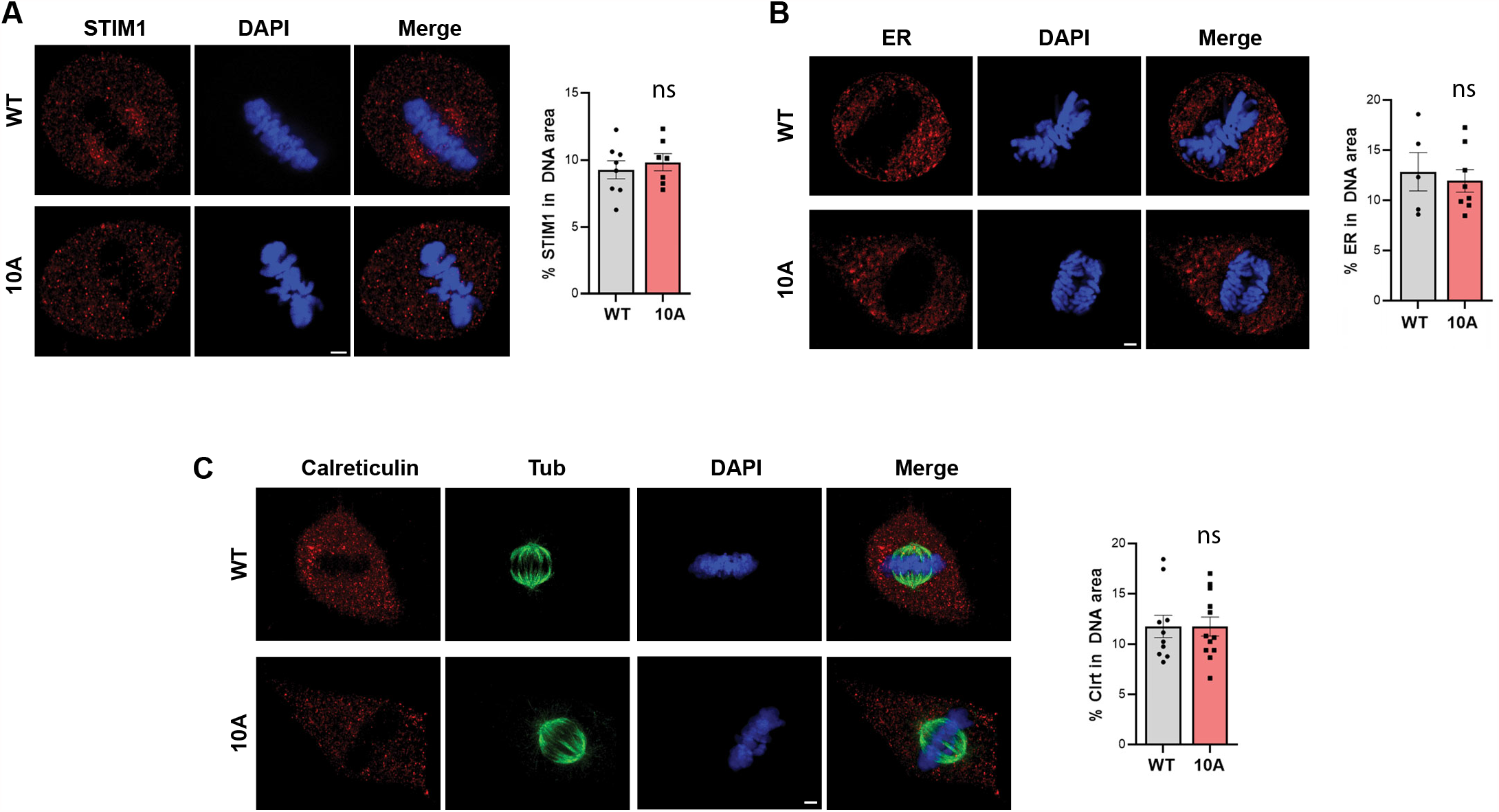
ER distribution is not disrupted in STIM1-10A MEFs. **(A-C)** Representative confocal images and statistical analysis of ER localization to the DNA area in WT and STIM1-10A MEF cells stained with DAPI to visualize the chromosomes, anti-STIM1 antibody (N-term) (A), or calreticulin and tubulin (C), or expressing mCherry-KDEL to label the ER (B). All cells imaged are from naturally occurring mitotic cells. Scale bar 3 µm. Summary data are represented as mean ± SEM with individual values represented. A: n=7-8 cells; B: n=5-8 cells; and C: n=10-12. For all three panels the means are not significantly different, unpaired t test.

To confirm these findings, we generated stable MEF cell lines that express WT or 10A mCherry-tagged STIM1. These lines were generated using a STIM1 null parental line that is a MEF line derived from STIM1-KO mice (Oh-Hora et al., 2008). These cell lines offer the advantage of consistent expression of mCh-STIM1 without having to worry about the differential expression from cell to cell when using transient transfection approaches. We did not observe any overt ER mis-localization in confocal imaging in cells co-stained for tubulin and DNA (Fig. 3A). Furthermore, STIM1 WT and 10A localize normally to the ER as shown by the reticular ER distribution in the magnified insets (Fig. 3B). To quantitatively assess ER colocalization with the spindle we generated 3D reconstructions of serial confocal z-stack images (Fig. 3C) and quantified STIM1 localization to the spindle area using Imaris software. The percent of STIM1 localizing to the spindle volume (Fig. 3D) or the DNA volume (Fig. 3E) in these analyses was similar in the STIM1 WT and 10A cell lines. We further assessed the Pearson Colocalization Coefficient between STIM1 and tubulin and observed no difference between the two cell lines (Fig. 3F).

**Figure 3.**
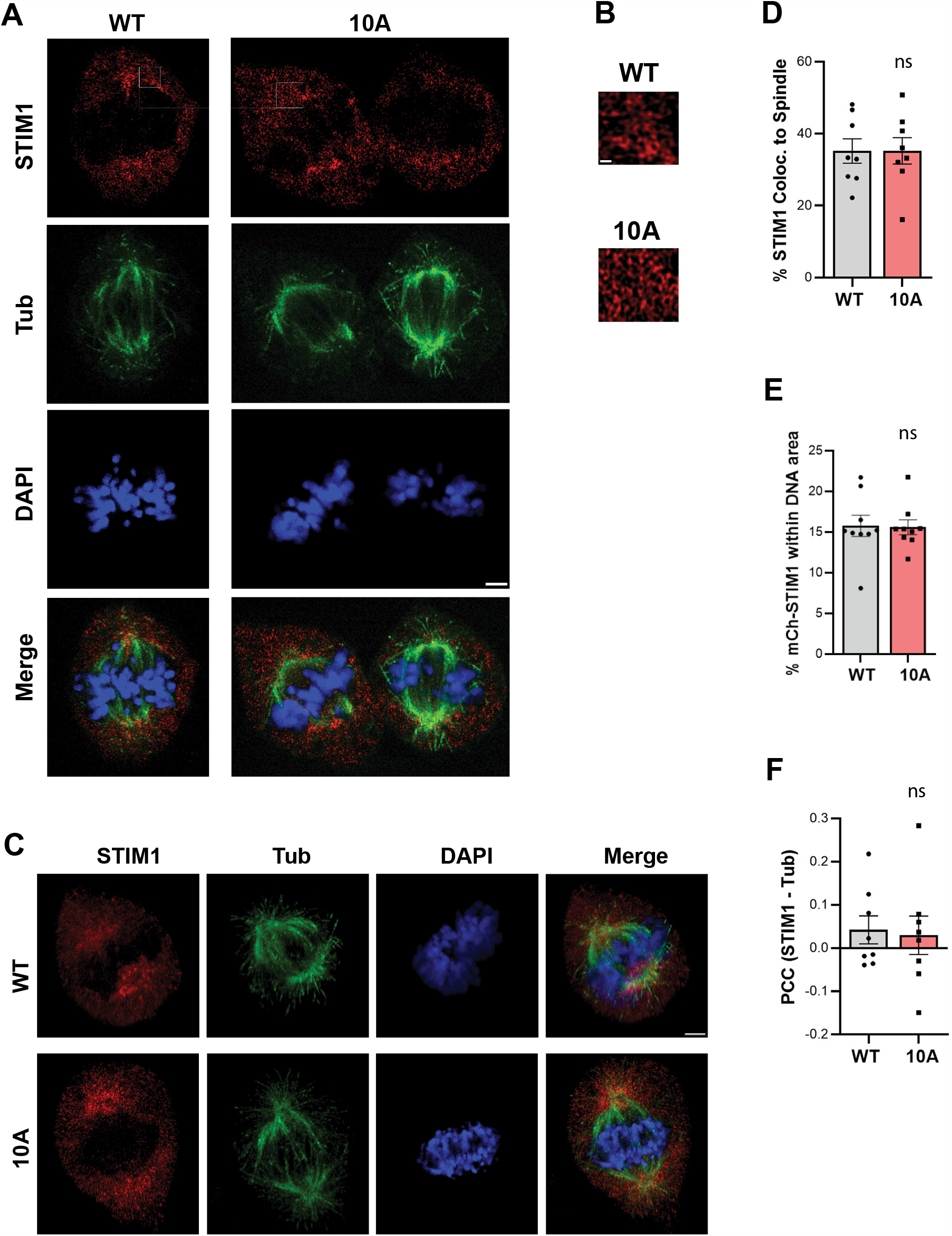
ER distribution is not disrupted in STIM1-KO MEFs stably expressing WT or 10A STIM1. **(A)** Representative images from STIM1-KO MEFs stably expressing wild-type STIM1 (WT) or the 10A mutant (10A) and stained for STIM1 (N-term antibody), tubulin and DAPI. **(B)** Magnification of the regions in panel A indicated by the white box to highlight the reticular ER structure revealed by STIM1 staining. Scale bar 3 µm. **(C)** Examples of 3D reconstructions of a confocal z-stack of images using Imaris software® to measure STIM1 colocalization in the 3D volume of the cell. **(D-E)** Percent of STIM1 localizing to the spindle (D) or DNA (E) volume occupied by the spindle. Mean ± SEM, n=8; ns, not significant, unpaired t test. **(F)** Pearson colocalization Coefficient between tubulin and mCh-STIM1 in the 3D volume of the cell calculated using the Imaris software colocalization tool. Mean ± SEM, n=8; ns, not significant, unpaired t test.

Taken together, our data show conclusively that phosphorylation of STIM1 during mitosis is not involved in ER partitioning since cells expressing only the 10A form of STIM1 distribute their ER normally without any overt defects. These results further argue that the previous observations were due to gross overexpression of STIM1-10A resulting in ER mis-localization, because even in cells stably expressing exogenous mCherry-STIM1 only (Fig. 3), we do not observe any ER mis-localization to the mitotic spindle.

### Role of STIM1 phosphorylation in cell migration

Previous studies have argued for an important role for STIM1 phosphorylation in cell migration (Casas-Rua et al., 2015; Lopez-Guerrero et al., 2020; Lopez-Guerrero et al., 2017). In particular, non-phosphorylatable STIM1 impaired cell migration, whereas constitutive phosphorylation promoted migration (Casas-Rua et al., 2015). Based on these findings one would expect that non-phosphorylatable STIM1-10A MEFs migration to be impaired. We therefore tested the cell migration potential of 10A MEFs as compared to their WT counterpart in three separate assays (Fig. 4). In the wound-healing assay we observe no statistically significant difference between 10A and WT MEFs in the rate of wound closure (Fig. 4A-B) or the speed of sheet migration (Fig. 4C). This is reflective of cell-sheet migration as cell proliferation is similar between STIM1-10A and WT MEFs (Fig. 1C). These results are consistent then with the similar SOCE levels observed in 10A and WT MEFs (Fig. 1E). Therefore, results from the wound healing assay argue against a role for STIM1 phosphorylation in cell-sheet migration.

**Figure 4.**
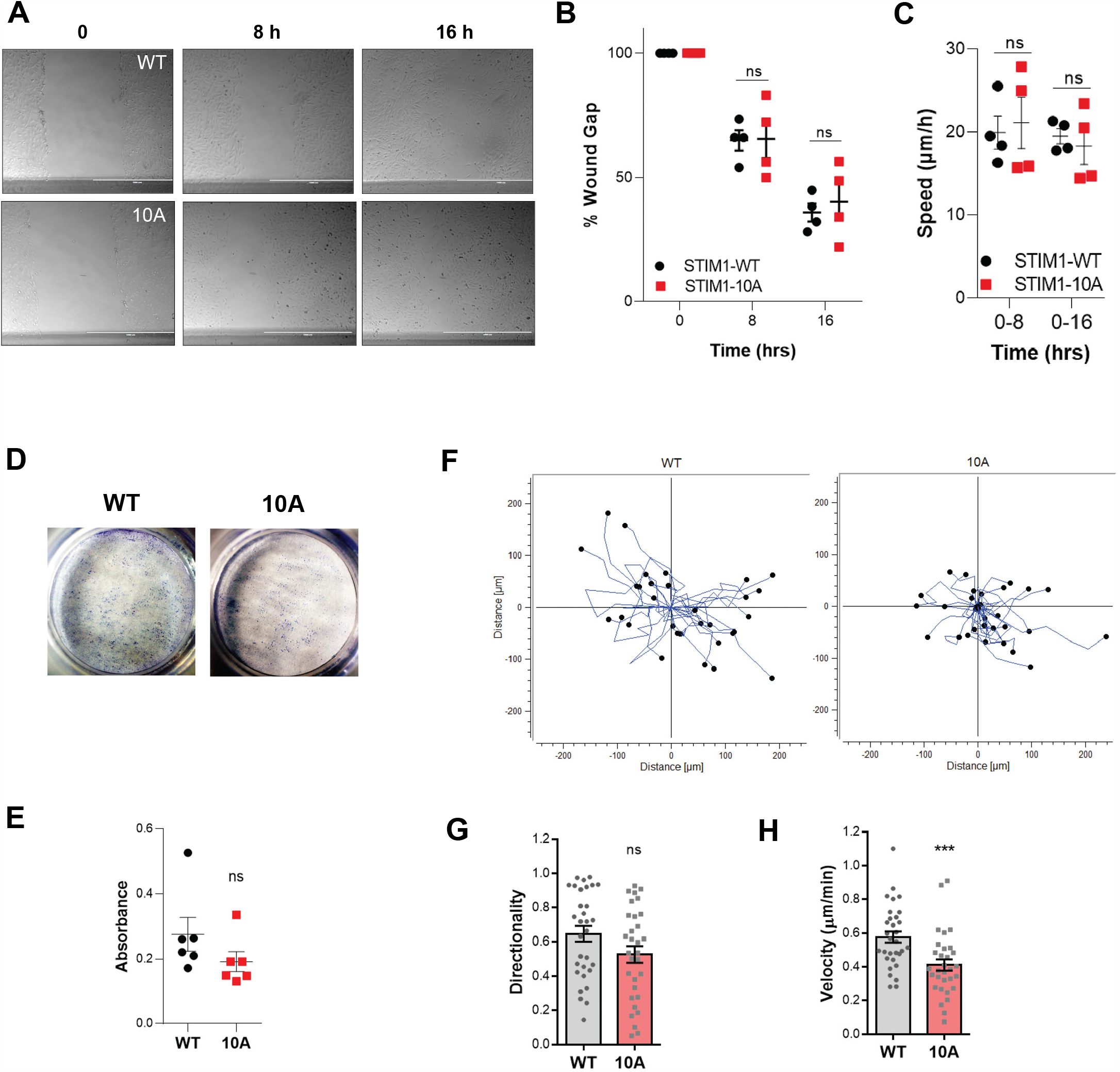
Role of STIM1 phosphorylation in cell migration. **(A)** Example images showing wound closure at 0, 8, and 16 hrs in WT and 10A MEFs. Scale bar 1 mm. **(B-C)** Summary data showing the rate of wound closure (B) and the speed of the migrating sheet of cells (C). Four independent experiments; mean ± SEM, ns, not significant, paired t test. **(D)** Representative images of transwell Boyden chamber invasion assays of STIM1-10A and WT MEFs stimulated with 10% v/v serum for 20 hrs. **(E)** Analysis of invasiveness by measuring cell density at a wavelength of 590 nm. n=6 wells from two independent experiments, mean absorbance ± SEM; ns, not significant, unpaired t test. **(F)** Overlapping live cell movement tracks from STIM1-WT and -10A MEFs. **(G-H)** Quantification of movement directionality (G) and cell velocity (H) in the two MEF lines. Mean ± SEM, n = 31 cells from three independent experiments; ns, not significant, *** p<0.001; unpaired t test.

Next, to test whether non-phosphorylatable STIM1-10A modulates cell invasion, we performed the transwell invasion assays with using Matrigel coated membranes. Matrigel is a complex mixture similar to basement membranes containing a small concentration of fibronectin. Cell invasion was induced by transferring transwells containing serum starved MEFs into complete media containing 10% FBS in the lower chamber. The results show no difference in the invasion capacity of 10A as compared to WT MEFs (Fig. 4D-E), arguing that STIM1 phosphorylation is not required for cell invasion.

Finally, to quantify the pattern and velocity of cell movement, we performed live cell tracking using epifluorescence microscopy. MEFs were grown in normal culture conditions on collagen I coated glass and time-lapse microscopy collected to generate individual cell tracks using manual tracking that were overlapped onto a single plot (Fig. 4F). We observed no difference in the directionality of individual cell migration between WT and 10A MEFs (Fig. 4G). In contrast, individual cell velocity was slightly but significantly lower in STIM1-10A MEFs as compared to WT (Fig. 4H). These results argue that in contrast to cell sheet migration in the wound healing assay, STIM1 phosphorylation may play a modulatory a role in individual cell migration.

### STIM1 phosphorylation state does not affect focal adhesions

Lead edge protrusion of migrating cells involves tightly coordinated changes in the actin cytoskeleton and the PM. The polymerizing actin filaments (F-actin) push the membrane outwards, forming the protrusive structures lamellipodia, filopodia and invadopodia (Hammad and Machaca, 2021; Ridley, 2011). To visualize actin filaments in live MEFs, we generated Ftractin-mCherry stable WT and STIM1-10A MEFs in which F-actin was labeled by a mCherry tagged actin-binding peptide derived from inositol 1,4,5-triphosphate 3-kinase A (ITPKA) (Hayer et al., 2016; Schell et al., 2001). The gross morphology of the actin cytoskeleton was similar in both cell lines as shown in the example cells in Figure 5A.

**Figure 5.**
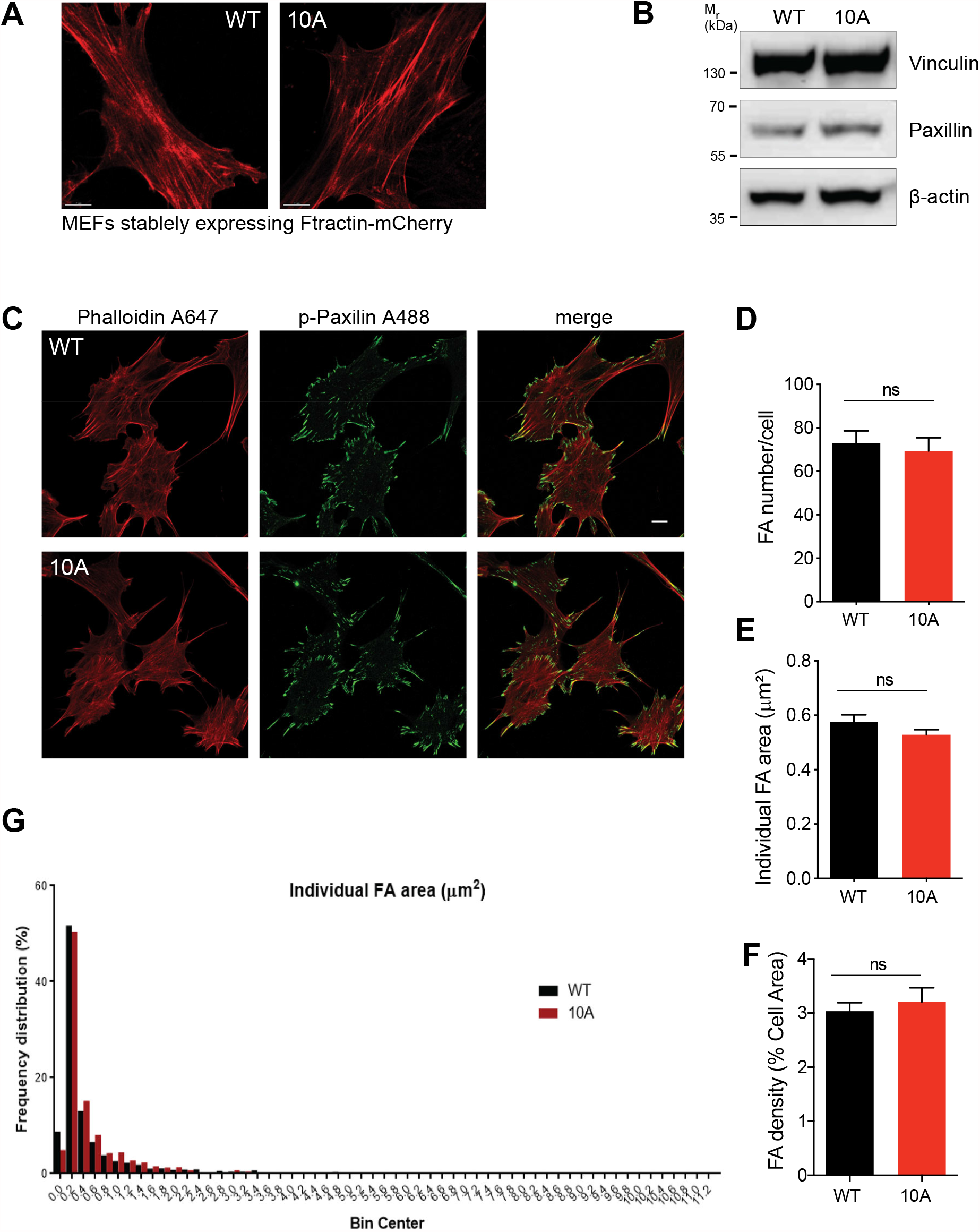
The STIM1 phosphorylation state does not alter focal adhesions. **(A)** Representative Airyscan confocal images of Ftractin-mCherry stable WT and STIM1-10A MEFs. Scale bar 7 μm. **(B)** Western blot of vinculin and paxillin in STIM1-10A and WT MEFs. Images represent two independent experiments. **(C)** Representative confocal images of focal adhesion (FA). MEFs were stained with anti-p-paxilin for FA and Phalloidin for F-actin. Scale bar 10 μm. **(D-G)** Quantification of FAs: number per cell (D), individual FA area based on p-Paxillin staining (E), focal adhesion density measured as the percent of cell footprint area occupied by FA (F), and the frequency distribution of individual FA area (G). For each group 20 cells from two independent experiments were analyzed. Mean ± SEM; ns, not significant, unpaired t test.

We next assessed focal adhesions (FA) in both cell lines because focal adhesion dynamics are critical for cell migration and are modulated in complex fashion by SOCE (Hammad and Machaca, 2021). We examined the expression of two FA components paxillin and vinculin, which play important roles in cell migration (Legerstee et al., 2019; Turner et al., 1990). Western blots revealed that paxillin and vinculin are expressed to similar levels in STIM1-10A and WT MEFs (Fig. 5B). FAs serve as points of traction and as signaling centers that link integrins to the actin cytoskeleton to control cell migration with paxillin as a major component of FA (Ridley et al., 2003; Romer et al., 2006). The phosphorylation of paxillin (p-paxillin) by FAK upon integrin activation modulates cell migration dynamically (Lopez-Colome et al., 2017). We therefore stained STIM1-10A and WT MEFs for actin and phospho-paxillin to quantify FAs (Fig. 5C-G). We observed no differences between the two cell lines in FA per cell (Fig. 5D), their individual area (Fig. 5E), or their density (Fig. 5F), which was assessed as the percent of cell area covered by FAs (Fig. S2C). Furthermore, FA size frequency distribution was not different between the two cell lines (Fig. 5G), arguing against modulation of particular FA subsets by STIM1 phosphorylation status. This was important to assess because mature FA (large) and nascent FA (small) modulate cell migration differentially at the front and rear of migration cells and are themselves differentially modulated by SOCE (Hammad and Machaca, 2021).

## DISCUSSION

STIM1 phosphorylation has been implicated in cell migration, regulation of ER distribution during mitosis, and SOCE activation (Casas-Rua et al., 2015; Lopez-Guerrero et al., 2020; Lopez-Guerrero et al., 2017; Pozo-Guisado et al., 2010; Smyth et al., 2012). STIM1 is a microtubule plus-end tracking proteins (+TIPs) with a ‘TRIP’ sequence (Fig. 1A, EB) that matches the ‘SxIP’ consensus binding site for EB1. Phosphorylation around the ‘SxIP’ motif in other +TIPs such as APC, MCAK, and CLASP2 abrogate EB1 interaction and MT tracking (Honnappa et al., 2009; Kumar et al., 2009). In agreement with these findings that phosphorylation of STIM1 inhibits its interaction with EB1 as assessed by immunoprecipitation (Casas-Rua et al., 2015; Pozo-Guisado et al., 2013). This dissociation was argued to release STIM1 from MT and allow it to move to junctions to support SOCE. In contrast to these finding, STIM1 was shown to maintain its ability to bind to EB1 after store depletion in its clustered activated state (Chang et al., 2018), and this mechanism restricts STIM1 targeting to ER-PM junctions and thus prevents Ca^2+^ overload (Chang et al., 2018). Consistently, knocking down EB1 leads to an increase in SOCE by around 20% (Chang et al., 2018).

Overexpression of STIM1 S575,608,621A did not support SOCE and an increase in STIM1 phosphorylation was detected following store depletion that also resulted in ERK1/2 phosphorylation (Pozo-Guisado et al., 2010). In this study, we use MEFs derived from a mouse strain where endogenous STIM1 has been replaced with a non-phosphorylatable STIM1 with all 10 S/T residues that match the ERK/CDK consensus mutated to Ala. The control MEF line expresses wild-type STIM1 and is derived from a congenic mouse line. We show that SOCE is activated to similar levels following store depletion in WT and 10A MEFs, arguing against a role for STIM1 phosphorylation in SOCE activation. This is consistent with previous reports showing that expression of STIM1-10A or 10E mutants where all these residues were mutated to either Ala or Glu supported normal SOCE (Smyth et al., 2012; Yu et al., 2019; Yu et al., 2009). Furthermore, CD4^+^ T cells isolated from WT and STIM1-10A mouse strains had similar levels of SOCE (Yu et al., 2019). Also, importantly STIM1-10A mice develop and reproduce normally with no obvious defects compared to their WT counterparts (Fig. S1A) (Yu et al., 2019). A potential explanation for these discrepancies is that the S5757,608,621A mutant behaves differently from the 10A mutant. This seems unlikely though as different Ala point mutants or double mutants did not have any significant impact on SOCE levels when overexpressed (Yu et al., 2009). Collectively, these findings argue against STIM1 phosphorylation at ERK/CDK sites playing a role in regulating SOCE amplitude following store depletion. They however do not rule out a possible role for other kinases in regulating SOCE. In that context, store depletion has been shown to induce STIM1 phosphorylation on Y361 through proline-rich kinase 2 (Pyk2), which in turn regulates STIM1-Orai1 interaction (Yazbeck et al., 2017).

The modulation of the STIM1-EB1 interaction by phosphorylation was also implicated in ER distribution during mitosis. Overexpressed STIM1 co-localizes with endogenous EB1 and tracks to MT in interphase but dissociates from MT in mitosis (Smyth et al., 2012). In contrast, STIM1-10A binds MT in mitosis and drags the ER with it to the mitotic spindle (Smyth et al., 2012). However, under these overexpression conditions a significant percent (∼40%) of STIM1-10A localizes away from the spindle, arguing that overexpression may be associated with mis-localization (Smyth et al., 2012). Here we show using multiple ER markers including STIM1 itself that the lack of endogenous STIM1 phosphorylation during mitosis does not alter ER distribution, and we observe no ER localization to the mitotic spindle. However as discussed above the evidence that STIM1 phosphorylation dissociates it from EB1 is compelling. Why then would STIM1-10A at endogenous expression levels not bind to MT and associate with the mitotic spindle in mitosis? We would argue that the ER remodeling observed in mitosis generates a stronger pull on STIM1 than that exerted by the STIM1-EB1 binding interaction. Although STIM1-10A in mitosis still has the ability to bind EB1, this interaction is not strong enough to counter the general ER remodeling that pulls the ER away from the spindle to not interfere with chromosome segregation and equal ER partitioning to the daughter cells. There is in fact direct evidence for such a mechanism in oocyte meiosis where the ER undergoes dramatic remodeling as well (Yu et al., 2009). In this case as expected expression of the constitutively active STIM1-D76A mutant localizes the ER-PM junctions and interacts with Orai1 in interphase, however in meiosis this interaction is disrupted and STIM1-D76A localizes exclusively to the remodeled ER away from Orai1 (Yu et al., 2009). As STIM1-D76A is constitutively active and binds Orai1, this shows that the dragging force on STIM1 exerted by ER remodeling overwhelms the STIM1-Orai1 interaction. The same would be expected for STIM1-EB1 interactions and ER remodeling in mitosis.

SOCE modulates Ca^2+^ signaling dynamics which are critical for cell movement (Hammad and Machaca, 2021). Furthermore, STIM1 phosphorylation has been implicated in cell migration (Casas-Rua et al., 2015; Lopez-Guerrero et al., 2020; Lopez-Guerrero et al., 2017). The STIM1 S575,608,621A mutant inhibits cell migration (Casas-Rua et al., 2015). Phospho-STIM1 is enriched preferentially at the leading edge of migrating cells and colocalizes with both Orai1 and the contractin, a regulator of membrane ruffling (Lopez-Guerrero et al., 2017). Finally, the Orai1-contractin interaction is dependent on the small GTPase Rac1 and regulates lamellipodia extension (Lopez-Guerrero et al., 2020). In contrast to these findings, we observe no differences between WT and 10A MEFs in cell-sheet migration and cell invasion in the wound healing and Matrigel invasion assays respectively. Furthermore, individual cell tracking reveals a slight decrease in cell velocity in 10A cells with no difference in directionality. This difference could be a reflection of the distinct matrices used in the three different assays. The wound healing assay was performed on plastic, the Transwell assay using Matrigel which is the closest to physiological ECM, and the individual cell movement on collagen I. The fact that we observe a decrease in velocity only in the individual cell tracking assay argues against an inherent role for STIM1 phosphorylation in cell migration but potentially for a role in regulating a particular subset of integrins that link to collagen I. In support of this conclusion, the STIM1-10A mouse line exhibits no overt developmental or other phenotypes and reproduces normally compared to WT animals (Yu et al., 2019). This argues that the decrease in cell migration velocity observed in 10A MEFs does not translate into a cell migration defect in the whole animal, which may be expected since the ECM in vivo is distinct from collagen I. Furthermore, we do not observe any alterations in focal adhesion size, density, distribution, or expression of molecular markers in STIM1-10A MEFs. This shows that the phosphorylation state of STIM1 does not alter FA dynamics.

Collectively our results argue against a prominent role for STIM1 phosphorylation in regulating cell migration. It is important to note though that the evidence supporting a role for SOCE in cell migration is compelling, where changes in the expression levels or function of STIM1 or Orai1 consistently affect cell migration in normal and cancer cells (Hammad and Machaca, 2021). This however does not seem to involve STIM1 phosphorylation at least at endogenous or near endogenous expression levels.

Collectively, we provide solid evidence that phosphrylation of STIM1 at ERK/CDK sites is dispensible for SOCE activation, ER distribution in mitosis, and cell migration. The findings from previous studies implicating STIM1 phosphorylation in these processes appear to be due to gross expression of STIM1 mutants which could be associated with mass action effects that would not otherwise apply to endogenous STIM1. This however leaves open for future studies the important question of the role of the dynamic STIM1 phosphorylation in cell physiology.

## METHODS

### Generation of STIM1-10A and WT MEF cell lines

The mouse embryonic fibroblast (MEF) cell lines were generated from the STIM1-10A mouse strain and the parental strain with a knocked-in floxed cassette that expresses wild-type STIM1 (WT). Both mouse strains were previously described (Yu et al., 2019). In this paper we refer to the MEFs from the parental line at WT for simplicity as STIM1 is normally phosphorylated in this line. However, it should be noted that given the genomic structure of parental line it eliminates some of the known STIM1 splice variants (Yu et al., 2019). Isolation of STIM1-10A and WT MEFs was performed according to standard methods. Briefly, the primary MEFs were prepared from E13.5 embryos. Embryo bodies were minced and digested with trypsin to isolate MEFs that were cultured in DMEM supplemented with 10% FBS (Gibco) and 1% penicillin and streptomycin. Primary MEFs were immortalized at passage 4 by infection with an SV40 T-antigen expressing recombinant lentiviral vector supernatant (Capitol Biosciences, CIP-0011) in the presence of 10μg/mL Polybrene at approximately 50% confluence for 24 hrs. Cells were continuously grown in the medium for 48-72 hrs after transduction to reach confluency. MEFs (passage 1) were sub-cultured into a 10 cm tissue culture dish. The MEFs were split every 3-4 days for 2 weeks after P1, then SV40 transformed clones were selected and plated for expansion. Clones were confirmed for SV40 transformation via PCR for SV40 T-antigen (Fig. S1B). The primer pair for SV40 is: 5′ TGAGGCTACTGCTGACTCTCAACA 3′ and 5′ GCATGACTCAAAAAACTTAGCAATTCTG 3′; the primer pair for GAPDH is: 5′ ACCACAGTCCATGCCATCAC 3′ and 5′ TCCACCACCCTGTTGCTGTA 3′.

### Cell culture and transfection

Cells including MEFs and STIM1-KO HEK293 cells (Emrich et al., 2019) were maintained in DMEM (Invitrogen) supplemented with 10% FBS and 1% Penicillin/Streptomycin (Invitrogen). Cells were maintained at 37°C in a 5% CO_2_-humidified incubator. To generate Ftractin-mCherry stable STIM1-10A and WT MEFs, cells were infected with lentivirus expressing Ftractin-mCherry (Hayer et al., 2016) (Addgene #85131). The plasmid pLenti-EB1-EGFP was obtained from Addgene (plasmid # 118084). To generate stable cell lines with the ER tagged, WT or STIM1-10A MEFs were infected with a retrovirus expressing mCherry-KDEL, which is retained in the ER. To generate MEFs expressing mCherry-STIM1 or mCherry-STIM1-10A, STIM1-KO MEFs were infected with a retrovirus expressing either mCherry-STIM1-10A or mCherry-STIM1. The DNA constructs of mCherry-STIM1-10A (S486A, S492A, S575A, S600A, S608A, S618A, S621A, T626A, S628A, S668A) and mCherry-STIM1 WT were described previously (Yu et al., 2019). Retroviruses expressing mCherry-KDEL, mCherry-STIM1-10A and WT STIM1 were constructed by inserting PCR fragments into XhoI-HpaI sites of pMSCV-puro vector (Clontech) and were packaged using the Phoenix-ECO cell line (ATCC CRL-3214). To obtain bone marrow derived macrophages (BMDM), bone marrows were cultured and *in vitro* differentiated in bone marrow differentiation medium containing 40 ng/ml recombinant murine M-CSF (PeproTech) as described before (Sun et al., 2015).

### Immunoprecipitation and Western blots

Cell lysates were separated on SDS-PAGE using NuPAGE 4-12% Bis-Tris Gels (Invitrogen). Proteins were transferred to PVDF membranes (Biorad), and membranes were washed, blocked, and incubated (12 hrs, 4 °C) with primary antibody in TBST (137 mM NaCl, 20 mM Tris, 0.1 % Tween-20, pH 7.6) supplemented with 3% (w/v) BSA (Sigma-Aldrich). PVDF membranes were washed and revealed using horseradish peroxidase (HRP)-conjugated secondary antibodies and ECL detection reagent (GE Healthcare). The blots were imaged on a Geliance gel imaging systems (PerkinElmer) with GeneSnap software. For immunoprecipitation (IP), lysates were incubated overnight at 4 °C with a STIM1 N-terminal polyclonal antibody with constant rotation, then pulled down with Protein A/G agarose beads (Santa Cruz) for an additional 1 hr at 4 °C with constant rotation.

Primary antibodies used are STIM1 polyclonal antibody (Proteintech #11565-1-AP, lot No.00016319, 1:100 for IP, detect N-terminal of STIM1), STIM1 polyclonal antibody (Cell Signaling Technology #4916, 1:1000, detect extreme C-terminal end of STIM1), STIM1 (D88E10) C-terminal rabbit monoclonal antibody (Cell Signaling Technology #5668, 1:1000, detect region around Pro622 of STIM1), EB1 (1A11/4) mouse monoclonal (Santa Cruz #sc-47704) and β-actin (C4) mouse monoclonal antibody (Santa Cruz #sc-47778). Paxillin (# ab2264, 1:1000) and vinculin (# ab129002, 1:1000) were purchased from Abcam. Both HRP-conjugated goat anti-rabbit-IgG and goat anti-mouse-IgG antibodies (1:5000) were purchased from Jackson ImmunoResearch Laboratories.

### Proliferation assay

STIM1-10A and WT MEFs were seeded at density of 5000 cells/well in 96-well microplate and incubated at 37 °C. Each group had 4 replicates along with control wells with only media to account for background fluorescence. Cell proliferation was performed by live cells staining with alamarBlue cell viability reagent (Invitrogen #DAL1100) at 6, 12 and 24 hrs and signal was detected by a CLARIOstar plus plate reader (BMG Labtech).

### Calcium imaging

STIM1-10A and WT MEFs were cultured on glass bottom 35 mm microplates until reaching 70-80 % confluency. Cells were loaded with 1 μM Fura-2 AM (Invitrogen #F1221) in medium and incubated for 25 min at 37°C. Cells were washed in Ca^2+^-free Ringer and then incubated in 1μM Thapsigargin (Invitrogen #T7459) to deplete Ca^2+^ store followed by addition of 2 mM Ca^2+^ to assess SOCE. Imaging was performed with alternating excitation at 340 nm and 380 nm while collecting emission at 510 nm using a PTI imaging system mounted on an Olympus IX71 with continuous perfusion. The analysis was performed on PTI EasyRatioPro1.6.0.101 where individual cells were selected (>20 cells per experiment) and the fluorescence readings were compiled and averaged. SOCE was calculated by subtracting fluorescence ratios (F340/F380) before Ca^2+^ addition from the highest value after restoration of extracellular Ca^2+^. Graphs were analyzed in Prism 8 software (GraphPad).

### Wound healing assay

STIM1-10A and WT MEFs were seeded at an equal density onto a 6-well plate and incubated at 37°C for 24 hrs. After incubation, the monolayer was scratched twice with a p200 pipette tip to form 2 lines on each well. Cell migration into the gap was recorded at 0, 8 and 16 hrs using an EVOS microscope. The data were collected from three independent experiments and the images were analyzed on Axiovision SE64 Rel 4.9.1 Imaging software where the gap on each image was measured by drawing 5 lines to calculate the average gap width. The percentage wound gap remaining and speed of migration were calculated and analyzed by using GraphPad Prism 8.

### Boyden chamber transwell invasion assay

MEFs (5×10^4^ cells) were plated in 8 µm pore-transwell BioCoat growth factor reduced matrigel invasion chambers (Corning, Catalog No. 354483) following the manufacturer’s instructions. Cell migration were induced by adding 10% v/v FBS in the lower chamber for 20 hrs at 37°C. Non-migrating cells were removed, and invading cells were fixed with formaldehyde (3.7%) and permeabilized with cold methanol (4°C) and then stained with 0.4% w/v crystal violet dissolved in 10% v/v ethanol. Membranes were washed with PBS and images collected using a ZEISS Stemi 508 stereomicroscope. Cell invasion was quantified colorimetrically using a CLARIOstar plus microplate reader. Briefly, 10% acetic acid was added and then incubate for 30 secs with shaking to lyse invading cells on the membrane and release the crystal violet. Optical density was quantified at 562 nm.

### Live cell tracking and confocal imaging

For live cell tracking, MEFs were seeded at low density and grown on 35-mm glass-bottom dishes (MatTek) coated with collagen I (Gibco #A1048301) at 37 °C with 5% CO2 for 24 h before imaging. Time lapse live cell brightfield imaging was performed at 37 °C on a Zeiss Z1 AxioObserver inverted fluorescence microscope using a 20x objective for 12 hrs at 1 image/3 min. Images were acquired using Zeiss Zen Blue software and ImageJ was used to manually track individual cell movements. Chemotaxis and Migration Tool software (Ibidi®) was used to plot cell tracks and compute velocity and directionality.

For confocal imaging, cells were imaged in Ringer solution containing 125 mM NaCl, 5 mM KCl, 1.5 mM CaCl_2_, 1.5 mM MgCl_2_, 10 mM D-glucose, and 20 mM HEPES, pH 7.4. Antibodies and reagent used for staining were STIM1 polyclonal antibody (Proteintech #11565-1-AP), α-Tubulin (DM1A) Mouse mAb (CST #3873), Calreticulin Antibody (CST #2891), Phospho-Paxillin (Tyr118) rabbit polyclonal antibody (CST #2541), SERCA2 ATPase Antibody (2A7-A1) (Novus #NB300-581), Alexa Fluor 488 conjugated Goat anti-Rabbit IgG antibody (Invitrogen #A11008), Alexa Fluor 488 conjugated Goat anti-Mouse IgG antibody (Invitrogen #A11029), Alexa Fluor 546 conjugated Goat anti-Rabbit IgG antibody (Invitrogen #A11035), Alexa Fluor 647 Phalloidin (Invitrogen #A22287), and Antifade Mounting Medium with DAPI (VECTASHIELD H-1200). Imaging was performed on a Zeiss LSM 880 confocal microscope with Airyscan using a Plan Apo 63x/1.4 oil DIC II objective with pinhole 1AU. Quantitative analysis of focal adhesion (FA) numbers, size and area was done using FIJI ImageJ analysis software from cells stained with anti-p-Paxillin and phalloidin (Fig. S2C). Images were processed with thresholding and FA numbers, size and area were quantified using the ‘Analyze Particles’ plugin in ImageJ.

### Statistics

Data are presented as mean ± SEM. Groups were compared using the Prism 8 software (GraphPad) using either paired or unpaired two-tailed Student t test as indicated. Statistical significance is indicated by p-values (>0.05 ns; <0.05 *; <0.01 **; <0.001 ***).

## AUTHOR CONTRIBUTIONS

ASH and WSB performed experiments and analyzed data. AE and EAA performed some experiments. FY performed experiments, analyzed data and wrote the first draft of the manuscript. KM analyzed data and wrote the final version of the manuscript.

## ACKNOWLEDGEMENTS

We are grateful to Mohamed Trebak (Pennsylvania State University) for providing the STIM1-KO HEK293 cell line, and Masatsugu Oh-hora (Tokyo Medical and Dental University) for the STIM1-KO MEF cell line. We thank the Microscopy Core at WCMQ for helping in experiments. The Core is supported by the ‘Biomedical Research Program at Weill Cornell Medical College in Qatar’, a program funded by Qatar Foundation. The statements made herein are solely the responsibility of the authors.

